# Neuronal ON/OFF Motion Detection Circuits Underlying Looming-Evoked Escape Behavior in *Drosophila*

**DOI:** 10.1101/2019.12.26.883587

**Authors:** Yeosun Kyung, Richard B. Dewell, Herman A. Dierick, Fabrizio Gabbiani

## Abstract

In *Drosophila*, early visual processing of motion information segregates in separate ON and OFF pathways. These pathways have been studied in the context of local directional motion detection leading to the encoding of optic flow that provides visual information for flight stabilization. Less is known about their role in detecting impending collision and generating escape behaviors. ‘Looming’, the simulated approach of an object at constant speed towards an animal, provides a powerful stimulus eliciting jump escape behaviors in stationary flies. We presented looming stimuli mimicking the approach of either a dark object on a bright background or a light object on a dark background, while inactivating neurons belonging either to the ON- or the OFF-motion detection pathways by expressing the dominant *Drosophila* temperature-sensitive mutant *shibire*^*ts*^ in different cells of the ON/OFF pathway. Inactivation of ON, respectively OFF, neurons led to selective decreases in escape behavior to light, resp. dark, looming stimuli. Quantitative analysis showed a nearly perfect splitting of these effects according to the ON/OFF type of the targeted neural populations. Our results suggest that *Drosophila* ON/OFF motion detection pathways play an important role in controlling jump escape responses according to looming stimulus polarity. They further imply that the biophysical circuits triggering *Drosophila* jump escape behaviors likely differ substantially from those characterized in other arthropods.

**Summary:** Inactivating fly neurons of the ON or OFF directional motion detection pathways during escape behavior selectively reduced jump responses to light and dark looming stimuli, respectively.

## Introduction

Visually-guided collision avoidance behaviors are critical for survival in many animals. They have been studied in a variety of species by using dark looming stimuli that simulate the approach of an object at constant speed towards the experimental subject (Fotowat and Gabbiani 2011; Peek and Card 2016; Schiff 1965). The ensuing nearly exponential increase in the angular size subtended at the retina by the simulated approaching object has long been recognized to provide a powerful cue eliciting escape (Gibson 1958).

In *Drosophila*, one neuron implicated in jump escape behavior to looming stimuli is the giant fiber (Allen et al. 2006). The giant fiber evokes a fast (short-mode, < 7 ms) escape response by triggering the extension of the middle legs and the elevation of the wings nearly simultaneously (von Reyn et al. 2014). As the resulting escape sequence consisting of jump and subsequent flight is often poorly coordinated (Card and Dickinson 2008; Fotowat et al. 2009), it is thought to be used as a last resort, for example in the case of particularly fast approaching threats (von Reyn et al. 2014).

Another class of neurons that has been implicated in visually-guided collision avoidance behavior are Foma1 neurons (de Vries and Clandinin 2012). In the optic lobe, Foma1 neurons are located in the lobula plate, which, together with the lobula, forms the third stage of visual processing in the optic lobe. Foma1 neurons respond to looming stimuli and trigger jump escape behavior when stimulated optogenetically, as do other optic lobe neurons (de Vries and Clandinin 2012; von Reyn et al. 2017; Wu et al. 2016). Their optogenetic stimulation results in jumps with generally longer latencies than those triggered by the giant fiber (long-mode, ≥ 7 ms), but it can also lead to seizure-like leg extensions, suggesting that the Foma1 line also targets ventral nerve cord neurons (von Reyn et al. 2017). Here, we took the opposite approach by acutely silencing Foma1 neurons using a *Foma1-GAL4* driver to express *UAS*-*shibire*^*ts*^, which inactivates synaptic transmission of targeted neurons in a dominant and temperature-dependent manner (Kitamoto 2001). Our results suggest that the class of lobula plate Foma1 neurons plays an important role in generating looming-evoked escape jumps in stationary flies.

Because of their dendritic arborizations in the lobula plate, Foma1 neurons are expected to receive inputs from two distinct ON and OFF directionally-selective motion pathways ending there (Fig. 1A; Maisak et al. 2013; Shinomiya et al. 2019). To study the contribution of these pathways to the generation of visually-guided collision avoidance, we systematically silenced ON and OFF neurons and compared their effect on escape behaviors generated by light and dark looming stimuli. By and large, we find that the segregation between ON and OFF neurons explains the generation of escape behaviors to light and dark looming stimuli, respectively. Thus, our results extend to looming-evoked jump escape behavior a dichotomy extensively documented in the context of optomotor responses (e.g., Ammer et al. 2015; Clark et al. 2011; Fisher et al. 2015; Joesch et al. 2010; Maisak et al. 2013; Serbe et al. 2016; Strother et al. 2017).

**Figure 1.**
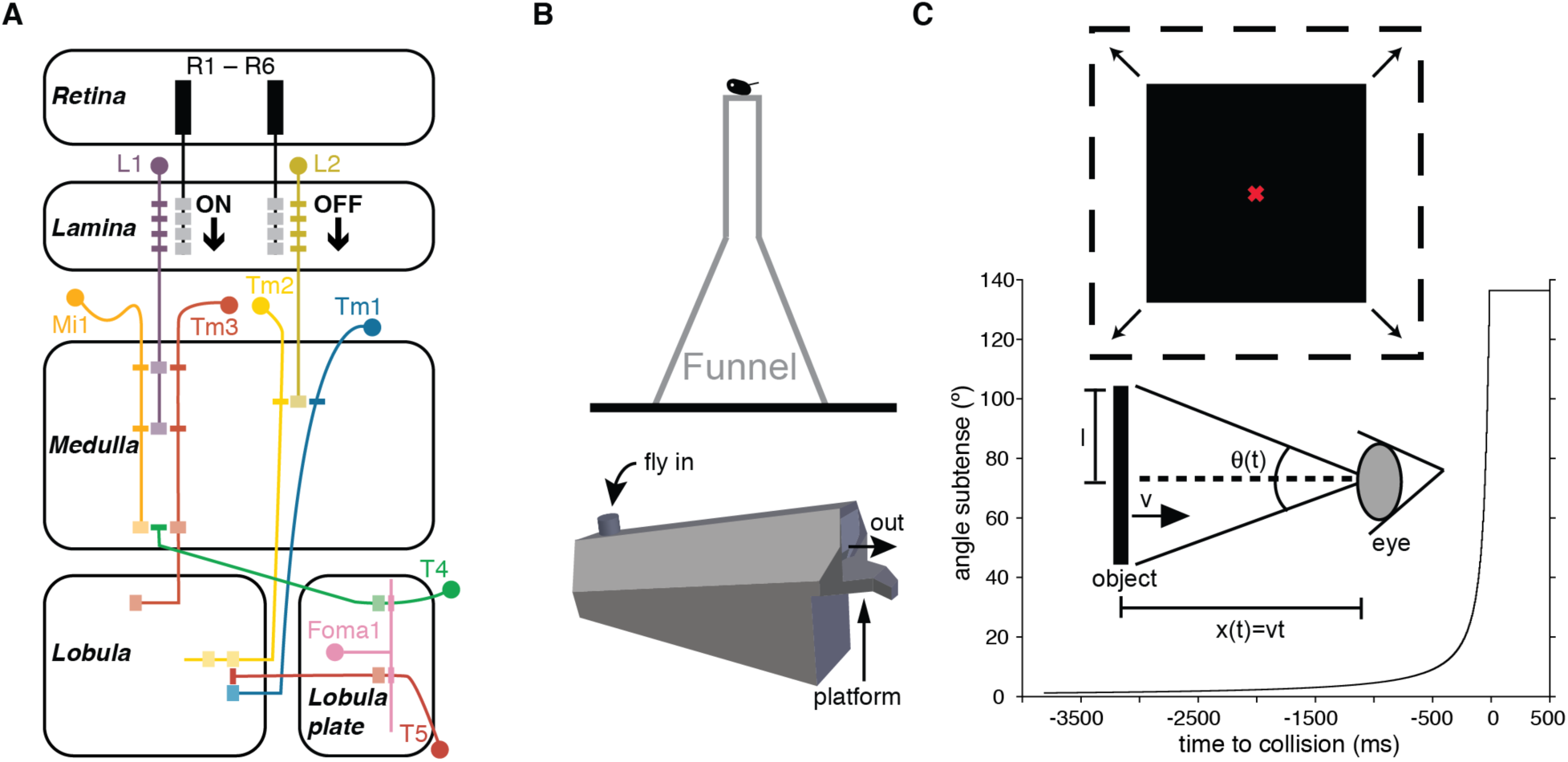
Neural pathways and behavioral setup. **A**: Main anatomical connections between the neurons studied here through *UAS-shibire*^*ts*^ inactivation. On the left are neurons thought to process ‘ON’ stimuli, including L1, Mi1, Tm3 and T4. On the right are those thought to process ‘OFF’ stimuli, including L2, Tm1, Tm2 and T5. Foma1 is thought to receives both T4 and T5 inputs. Presynaptic connections are indicated by thick bars in half tone; postsynaptic connections by thin bars in full tone. The same color code for the different cell classes is used in Figs. 2-4. **B**: Behavioral setup. Top, the fly was placed in an inverted funnel and was stimulated visually as it exited the top of the funnel. Alternatively, the fly was placed in a semi-transparent holder and was stimulated immediately after exiting it through a platform (bottom). **C**: Looming stimulus. The inverted funnel was placed in front of a screen so that the fly was approximately aligned with its center (top, red cross). The looming stimulus expanded symmetrically as indicated on top for a black square on a white background. Looming stimuli of inverted polarity were used as well. Alternatively, the semi-transparent holder was centered with its long axis parallel to the screen so that the right eye of the fly faced the screen upon exiting onto the platform. Bottom, time-course of the angle subtended by the expanding square relative to projected collision time. This stimulus simulates the approach of a black square of half-size, *l*, approaching towards the animal at a constant speed, *v* (middle).

## Materials and methods

### Fly lines

We used the following twelve lines, which target the neuronal populations indicated in parentheses. *Foma1-GAL4/+* (Foma1 in the lobula plate, de Vries and Clandinin 2012; also labels unidentified neurons in the ventral nerve cord, a gift from Tom Clandinin), *R42F06-GAL4/+* (BL# 54203, T4 and T5, Jenett et al. 2012; Maisak et al. 2013), *R54A03-GAL4/+* (BL# 50457, T4, Jenett et al. 2012; Maisak et al. 2013; Shinomiya et al. 2014), *R42H07-GAL4/+* (BL# 50172, T5, Behnia et al. 2014; Jenett et al. 2012; Shinomiya et al. 2014), *c202a-GAL4/+* (L1, Rister et al. 2007, a gift from Tom Clandinin), *21D-GAL4/+* (L2, Rister et al. 2007, a gift from Tom Clandinin; also labels Pm1 and Pm2 in the medulla), *NP6298-GAL4/+* (L1 and L2, Rister et al. 2007), *27b-GAL4/+* (Tm1, Behnia et al. 2014; Erclik et al. 2017; Morante and Desplan 2008, a gift from Tom Clandinin), *Bsh-GAL4/+* (Mi1, Behnia et al. 2014; Hasegawa et al. 2011, a gift from Tom Clandinin; also labels L4 and L5 in the lamina), *686-GAL4/+* (Mi1, Behnia et al. 2014, a gift from Tom Clandinin; also sparsely labels Tm2 in the medulla), *otd-GAL4/+* (Tm2, Morante and Desplan 2008, a gift from Tom Clandinin; also labels photoreceptors), *R13E12-GAL4/+* (BL# 49250, Tm3, Behnia et al. 2014; Jenett et al. 2012; also labels unidentified tangential cells in the medulla). The main documented anatomical connections between the neurons targeted by these lines are sketched in Fig. 1A. These lines were crossed with the line *UAS-shi*^*ts*^*/UAS-shi*^*ts*^ (located on the 3^rd^ chromosome, BL# 44222) carrying the mutant gene *shibire*^*ts*^ which rapidly abolishes synaptic transmission at elevated (permissive) temperatures (≥ 28.5 °C) in neurons expressing it through activation by GAL4 of the upstream activating sequence (UAS) promoter (Kitamoto 2001). Carrying out experiments at non-permissive (≤ 22.5 °C) and permissive temperatures for *shibire*^*ts*^ allowed us to test the effect of inactivating the targeted neuronal populations on jump escape behavior. Since the line *Foma1-GAL4/+* also expresses GAL4 in ventral nerve cord neurons, we generated a *tsh-GAL80; UAS-shi*^*ts*^ strain and crossed it to the *Foma1-GAL4* strain to test the effect of inactivating Foma1 neurons in the central brain. The *tsh-GAL80* line (generated by Julie Simpson, a gift from Gero Miesenböck) expresses GAL80 in the thorax, including ventral nerve cord neurons, thereby suppressing GAL4 expression there (Caygill and Brand 2016; Clyne and Miesenböck 2008; Röder et al. 1992; Suster et al. 2004). All lines were crossed with the fluorescent marker line *UAS-GFP/UAS-GFP* (JFRC81, Pfeiffer et al. 2012, a gift from Koen Venken) for immunolabeling and anatomical characterization of neurons expressing GAL4. Representative staining patterns for all lines are illustrated in Fig. 2B, and in Supp. Figs. 1, 3 (see also Supplementary Movies 1-13).

**Figure 2.**
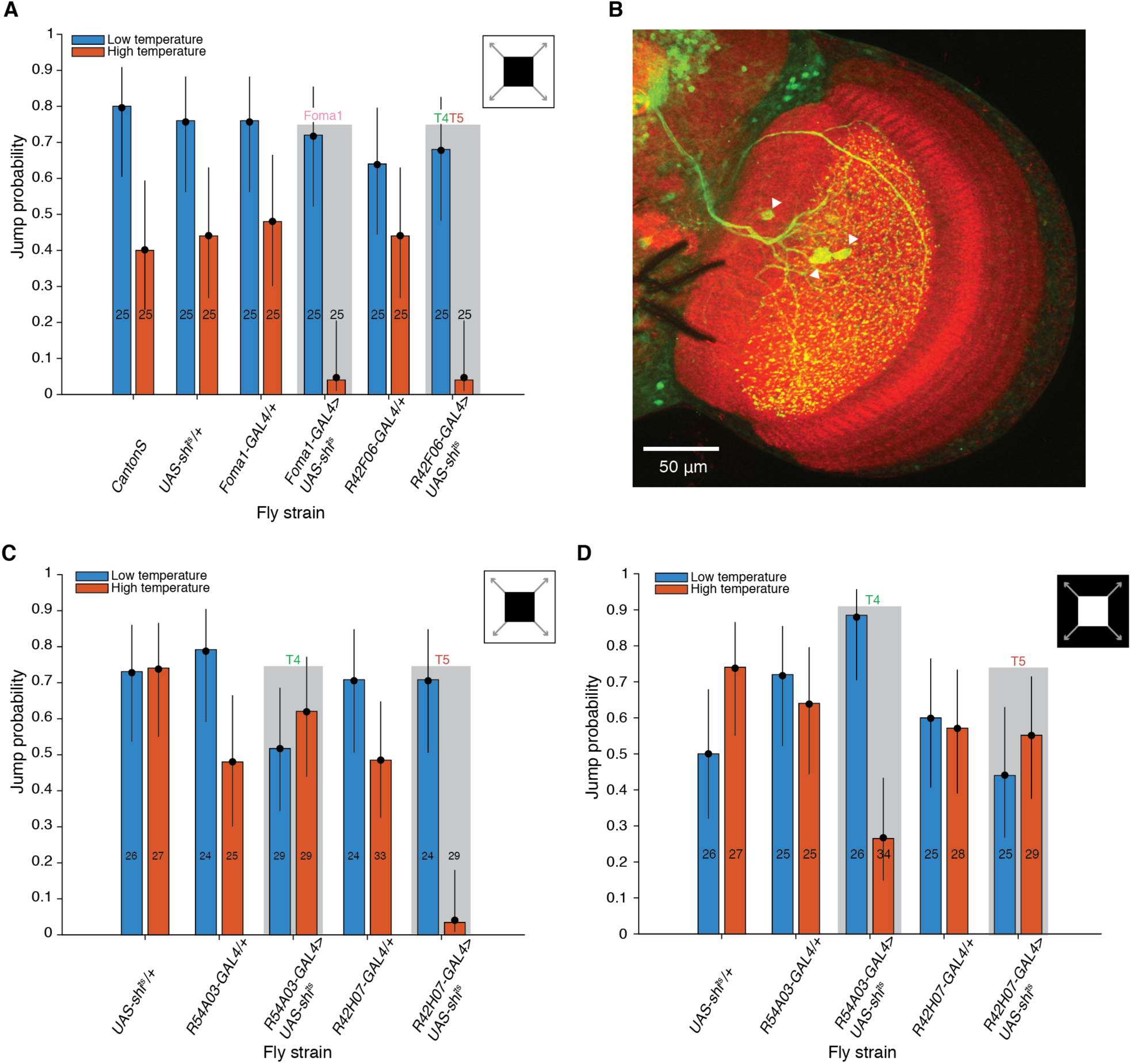
Effect of blocking lobula plate projecting neurons on jump escape behavior compared to Foma1 neurons. **A**: Jump probability of fly lines with either Foma1 or T4 and T5 neurons inactivated through *GAL4*>*UAS-shi*^*ts*^ at high temperature compared to low temperature (inside grey rectangles). Four control fly lines are illustrated as well on a *w*^*+*^ background. **B**: Immuno-staining of Foma1 neurons in the lobula plate. Three cell bodies are visible (arrowheads) as well as extensive dendritic arbors covering the entire lobula plate. Red: neuropil stain. **C, D**: Effect of inactivating either T4 or T5 neurons on jump probability to ‘ON’ and ‘OFF’ looming stimuli (inside grey rectangles), as well as in three associated control fly lines (*w*^*+*^ background). In A, C and D: total trial numbers reported in or on top of each bar. Additionally, dots indicate MUEs and vertical black lines 95% confidence intervals. Cell names are color-coded as in Fig. 1. Stimulus polarity indicated in top right insets.

### Immunostaining

Adult *Drosophila melanogaster* specimen were anesthetized on ice before dissection. After dissection, the brains were fixed in freshly made 4% paraformaldehyde (PFA) for one hour. The tissues were washed six times for 20 minutes using 1X phosphate buffer solution (PBS, 0.01 M PO_4_^3-^) containing 0.5% Triton X-100 to promote penetration of the antibodies. Goat serum (5%) was then added to block unspecific antigens. Antibodies were diluted in the same solution. We used the following primary antibodies: rabbit anti-GFP (1:500, A11122, Invitrogen, Carlsbad, CA, USA), and mouse monoclonal anti-Dlg (1:100, 4F3, Developmental Studies Hybridoma Bank). The primary antibodies were incubated with the tissues for 48 hours at the dilutions indicated above followed by regular washing (six times, as above). Chicken anti-rabbit tagged with Alexa Fluor 488 and goat anti-mouse tagged with Alexa Fluor 568 were used as secondary antibody at a dilution of 1:1000 (Invitrogen). Tissues were incubated with these secondary antibodies for 48 hours at 4 °C followed by regular washing. To remove unspecific binding as completely as possible, the tissues were washed for another 48 hours. Finally, the tissues were mounted in Slow Fade antifade reagent mounting media (Molecular Probes, Invitrogen) on a glass slide, cover-slipped and sealed with nail polish as previously described (Gnerer et al. 2015).

### Imaging

The brains and the optic lobes were imaged under an upright microscope with a 20X or 40X objective using a structured illumination module (Zeiss ApoTome and ZEN software, White Plains, NY, USA). The images shown in Fig. 2B and Suppl. Fig. 1 are maximum intensity projections from ∼4 to ∼12 slices through the optic lobe, or ∼30 slices through the ventral nerve cord of the animal. Optic lobe movies of the same lines were assembled from stacks of images acquired at the following pixel resolutions (x, y and z): 0.163 × 0.163 × 0.600 *µ*m (Supp. Movie 1 and 2, 40X) or 0.317 × 0.317 × 0.951 *µ*m (Supp. Movie 3-12, 20X).

### Behavior

Experiments were carried out in a small room (1.6 m^2^) containing the behavioral stage and three computers for stimulus presentation and data acquisition. During all experiments the fly strains were assigned a coded key unknown to the experimenter and revealed only after the experiment and basic data analysis were completed. This blinding procedure prevented experimental bias. Temperature was controlled manually using the room thermostat and a small electric heater. Temperature and humidity were measured with a hygro-thermometer positioned 15 cm away from the behavioral setup (Extech 44703, Nashua, NH). In most experiments, flies were placed in an inverted funnel similar to that described in (Fotowat et al. 2009). They were allowed to exit the funnel from its narrow tube at the top (Fig. 1B, top). In the experiments depicted in Fig. 4 we used a custom engineered semi-transparent fly holder that allowed higher throughput and more reliable positioning of the flies relative to the stimulation screen (described below). Flies were placed inside 10-20 such identical holders (Fig. 1B, bottom). They were allowed to adapt to this environment for at least ten minutes before placing one holder parallel to the stimulation screen and opening its front aperture, allowing the fly to exit onto a narrow platform with its right eye directly facing the stimulation screen. Shortly after flies exited the funnel or fly holder, a looming stimulus was manually activated (Fig. 1C). The stimulation screen was a cathode ray tube monitor refreshed at 200 frames per second and placed at 60 mm from the funnel or fly holder. The looming stimuli had a half-size to speed ratio, *l/*|*v*|, equal to 40 ms (*l*, half-size; *v*, approach speed, < 0 by convention so that time, *t*, before collision is < 0; Gabbiani et al. 1999). The stimuli simulated the approach at constant speed of a solid square either black or white on a white, resp. black background. The luminance of the screen at the black and white levels used was 2.5 and 72 cd/m^2^, respectively. Presentation of black or white stimuli occurred in pseudo-random manner over blocks of 20 stimuli (ten white and ten black). Stimulus presentation lasted 3.8 s. The stimulus subtended 1.2° at the start and 136.4° at the end of stimulation. During each trial, the temperature and humidity were recorded (Supp. Fig. 4) as well as the eventual occurrence of a jump. The position of the fly was also recorded when using the semi-transparent holders since some flies climbed on top of the holder instead of remaining on the platform. The behavior of each fly was filmed at 200 frames per second, in synchrony with each frame of the stimulus using a CCD camera (Gazelle, FLIR Systems Inc., Wilsonville, OR, USA), a 50 mm objective (Nikkor f/1.2, Nikon, Tokyo, Japan) and a 12 mm extension tube (Vello, EXT-ND, Gradus Group LLC, New York, NY, USA) mounted on the camera using a F to C mount adapter (Bower, C for Nikon, S. Bower Inc., Long Island City, NY). Infrared lightening was provided by six LEDs (S75, Smart Vision Lights, Muskegon, MI). Synchronization was achieved through a transistor-transistor logic (TTL) pulse issued by the video stimulation computer through its parallel port to the recording camera.

**Figure 3.**
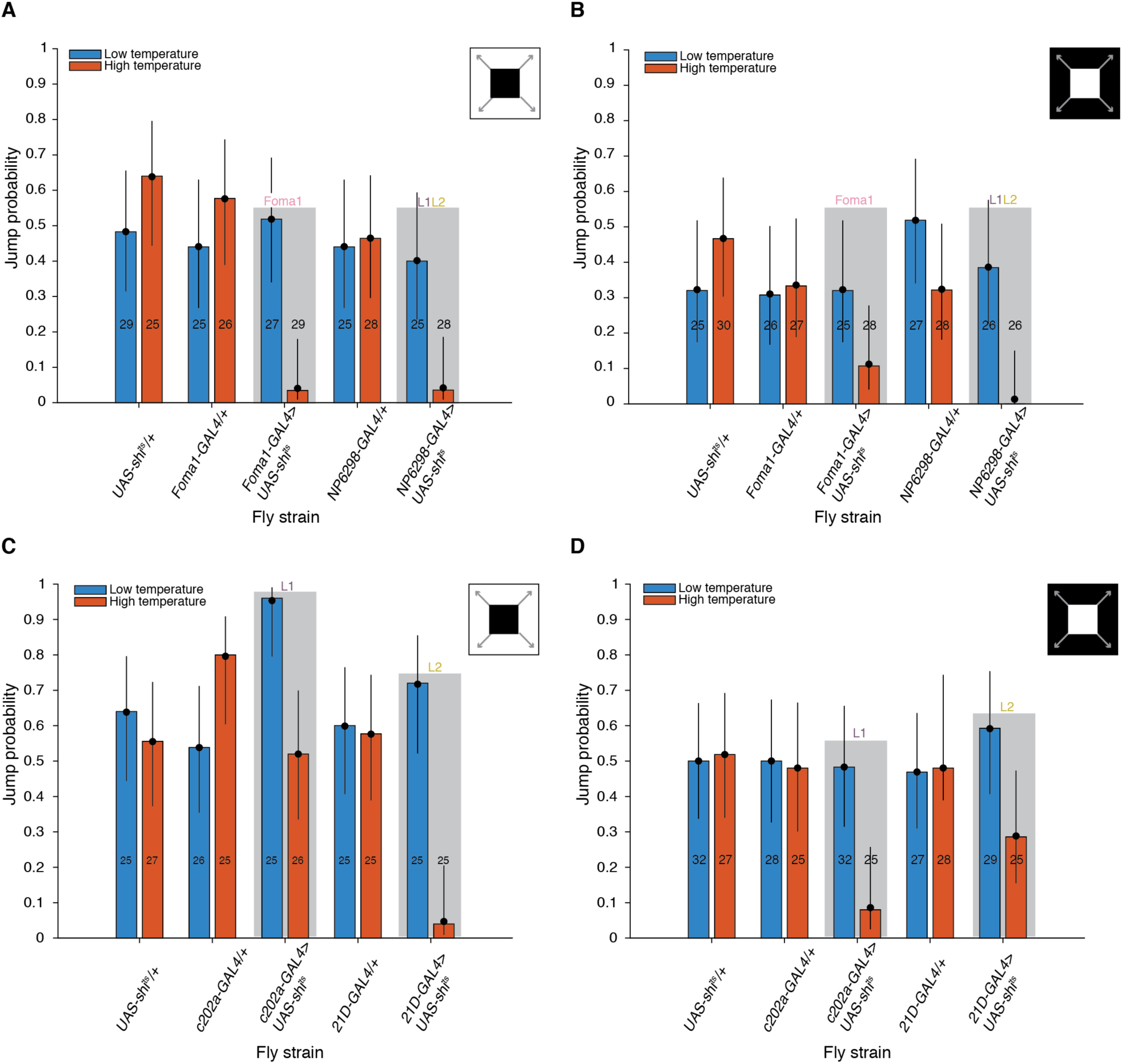
Effect of blocking lamina neurons on jump escape behavior compared to Foma1 neurons. **A, B**: Jump probability of fly lines with either Foma1 or L1 and L2 neurons inactivated through *GAL4*>*UAS-shi*^*ts*^ at high temperature compared with low temperature (inside grey rectangles; ‘ON’ and ‘OFF’ looming stimuli, respectively). Three control fly lines are illustrated on a *w*^*+*^ background. **C, D**: Effect of inactivating either L1 or L2 on jump probability to ‘ON’ and ‘OFF’ looming stimuli. Plotting conventions as in Fig. 2.

**Figure 4.**
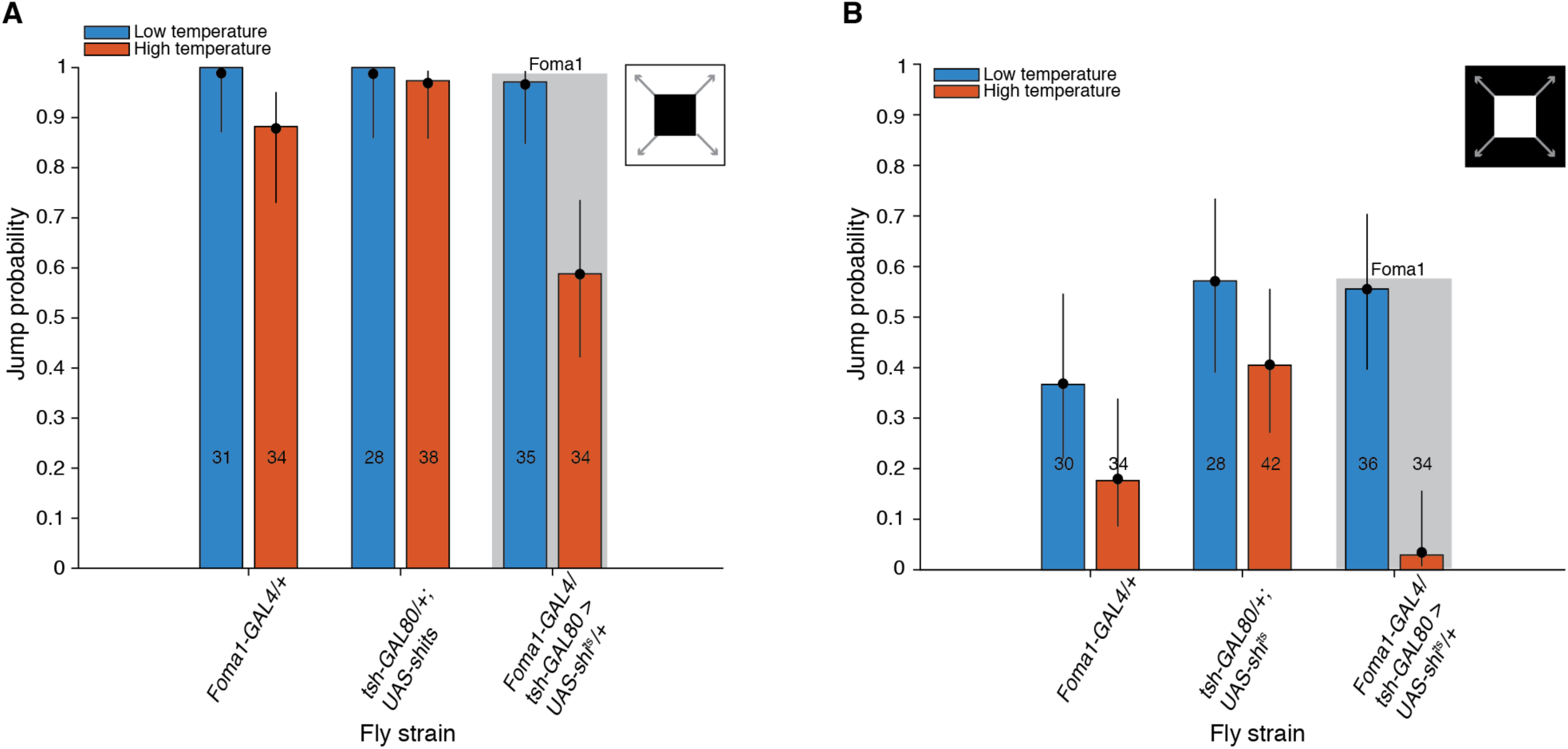
Behavioral effect of inactivating Foma1 mainly outside of the ventral nerve cord. **A, B**: Jump probability of the double mutant fly line *Foma1-GAL4/tsh-GAL80*>*UAS-shi*^*ts*^ whose *shibire*^*ts*^ expression is suppressed within the ventral nerve cord and thus inactivating Foma1 neurons mainly in the central brain at high temperature compared with low temperature (inside grey rectangles; ‘ON’ and ‘OFF’ looming stimuli, respectively). Two control fly lines are on a *w*^*+*^ background.

### Behavioral data analysis

#### Jump probabilities

Following each experiment, the video recording of each jump trial was inspected. When the occurrence of a jump was confirmed, the time (video frame) of the jump was recorded. A trial was invalid and discarded if the fly escaped within the first half second of stimulation as it was difficult to ascertain that such escape behaviors were triggered by the stimulus. Trials were also invalid if the fly’s head was oriented more than 135° away from a reference position facing the screen. In total we carried out eight behavioral experiments yielding 126 data sets and 3,370 valid trials. Jump probability was computed separately for each data set, corresponding to a fly strain, temperature range (high or low) and looming stimulus type (dark or light). The median number of trials per data set was 26 (minimum: 24, maximum: 42).

#### Jump preparation times

In *UAS-shi*^*ts*^*/+* flies, we had a sufficiently high number of trials to estimate the distribution of times used by flies to prepare for jump escape behavior. We extracted from each video of a jump the first frame of wing-raise. Preparation time was defined as the time from initial wing raise to the jump. The distribution of preparation times was binned at a resolution (5 ms) corresponding to the video frame rate (200 Hz). The fraction of short mode escapes (< 7 ms) was estimated as the fraction of preparations times in the 0-5 ms bin added to 2/5 of the fraction of preparation times in the 5-10 ms bin.

### Statistics

#### Power analysis estimation based on experimental sample size

For each data set, we computed the jump probability based on the number of jumps observed and the total number of trials. When testing if two jump probabilities differ, two types of error can be made. The first one is to declare the probabilities different when in fact they are identical. This is called a type I error and its probability is denoted by *α*. The second, type II, error is to declare the probabilities identical when they are different. Its probability is denoted by *β*. Equivalently, the probability 1 − *β* of correctly declaring them different, is called the power of the test. To distinguish an initial jump probability of 0.70 from a decrease to 0.25 with *α* = 0.05 and 1 − *β* = 0.80 requires 22 independent jump trials (Chap. 4 and Table A.4 of Fleiss et al. 2003). Our experiments were thus designed to use on the order of 25 jump trials per data set.

#### Median unbiased estimator of jump probability

If *n* independent jump trials are carried out with *x* positive outcomes, the maximum likelihood (ML) estimate of the binomial parameter corresponding to the jump probability is 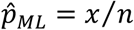. Because in our experiments the number of trials was relatively small (∼25), we also computed a robust estimator of jump probability, the median unbiased estimator (MUE, Hirji et al. 1989). By definition, a given estimator 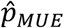 of the jump probability *p* is median unbiased if

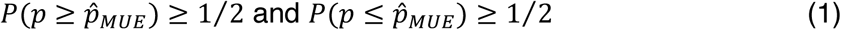

meaning that the true probability *p* has equal chance of being located on either side of 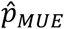. Due to the discrete nature of the data, there exists a range of estimators satisfying these conditions and one takes their midpoint as the MUE. For each experiment, 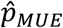 was computed from *x* and *n* following the method presented in sect. 2 of (Lin et al. 2009). For 0 < *x* < *n*, the lower and upper bounds, 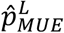 and 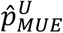, of the range of estimators satisfying Eq. (1) are given by

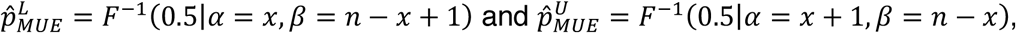

where *F*^−1^(0.5|*α, β*) is the 2-quantile (median) of the beta distribution with parameters *α* and *β*. Their midpoint is the desired MUE, given by

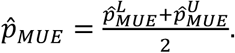

For *x* = 0 or *n*,

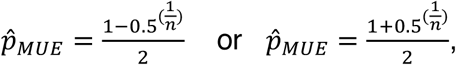

respectively. The MUE of jump probability was always closer to 0.5 than the ML estimator and thus more conservative (Results). In all jump probability plots the ML estimator is displayed as the height of a bar for each strain and the MUE by a dot.

#### Wilson score confidence interval

It has been recognized for some time that the confidence interval for proportions based on the normal approximation, called the Wald confidence interval, has poor statistical properties (Brown et al. 2001). The same is true for conventional bootstrap confidence intervals (Wang 2013). Instead, we computed the Wilson score interval, which like the MUE has been recommended based on extensive considerations (Brown et al. 2001; Wang and Hutson 2013). The Wilson score was calculated by inverting eq. (2.16) of (Fleiss et al. 2003), neglecting the continuity correction term (1/2*n*). Specifically, let Φ(*z*) be the normal cumulative distribution function and *z*_*α*/2_ = Φ^−1^ (1 − *α*/2) the critical value of the normal distribution for the desired two-tailed significance level *α* (in our case, set to 0.05). If *n* is the number of trials, 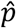 the estimated jump probability, then the upper and lower bounds, *w*_*U*_ and *w*_*L*_, of the Wilson score interval are given by

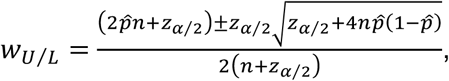

respectively.

#### Logistic regression

The MUE of jump probability and its 95% Wilson score confidence interval determine jointly the range where we expect the true jump probability to lie, and thus when two jump probabilities are highly likely to be different. In addition, we used logistic regression to quantify how strong the effect of temperature on the jump probability of a given fly strain was. This measure is independent of whether the jump probabilities at the two considered temperatures are significantly different or not based on the MUE of jump probability and the Wilson score interval. It allows to quantify in a graded manner subtler effects even when jump probabilities are not significantly different with high confidence. We illustrate the use of logistic regression in the case of a single fly strain with (estimated) jump probabilities *p*_*i*_, *i* = 1, 2, at low and high temperature, respectively. For further details, see McCullagh and Nelder (1989); Rodriguez (2007). In logistic regression one fits a linear model to the log-odds

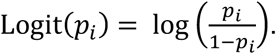

The simplest model assumes Logit(*p*_*i*_) = *η*, independent of *i*. A measure of discrepancy between the observed and fitted values is given by the deviance,

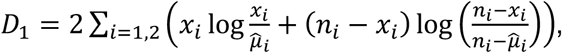

where *n*_*i*_ is the number of trials for condition *i, x*_*i*_ is the number of successes (jumps), and 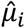 is the fitted number of successes, obtained by inverting the logit equation, *p*_*i*_ = e^*η*^ /(1 + e^*η*^) and multiplying by *n*_*i*_. By definition, the deviance is twice the product of the observed outcomes multiplied by the logarithm of the ratio of observed to predicted outcomes. Hence, if the fit is perfect the ratio of observed to predicted outcomes is one, their logarithm is zero and the deviance is equal to zero.

This last condition occurs in a saturated model, in which as many parameters as observations are fitted to the data. In the above example, such a model has two parameters,

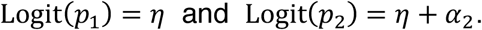

Its deviance, *D*_2_ = 0. In general, for such nested models the difference in deviance, *D*_1_ − *D*_2_, is asymptotically distributed as a *χ*^2^ random variable with *k* degrees of freedom, where *k* is the difference in the number of parameters of the two models, here *k* = 1. The difference in deviance provides a statistical test of the hypothesis *α*_2_ = 0. If *α*_2_ is different from zero, the fitting procedure yields its estimated value 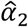 and the Wald confidence interval, 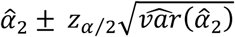, where 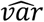 denotes the standard error of the estimate. By inverting once again the logit equation, we see that the odds are given by

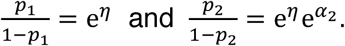

The odds ratio, 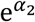, for the two populations provides a measure of the strength of the treatment. It is equal to one when the null hypothesis is satisfied (*α*_2_ = 0), and it is smaller (resp. greater) than one if the treatment causes a decrease (resp. increase) in the odds of the experimental outcome. In our case, it quantifies the relative change in the odds of jumping caused by the change in temperature. Its estimated value and confidence interval (*CI*) are obtained by exponentiating the corresponding values for *α*_2_. The values and standard error estimates of the log-odds ratios in Fig. 6 were computed similarly. Note that this procedure can be straightforwardly extended to multiple strains by using a single parameter *η* to fit the logit of all strains at low temperature and a second parameter *η* + *α*_2_ to fit the same logits at high temperature. Additionally, we used it to quantify changes in the odds of jumping for dark vs. light looming stimuli.

**Figure 5.**
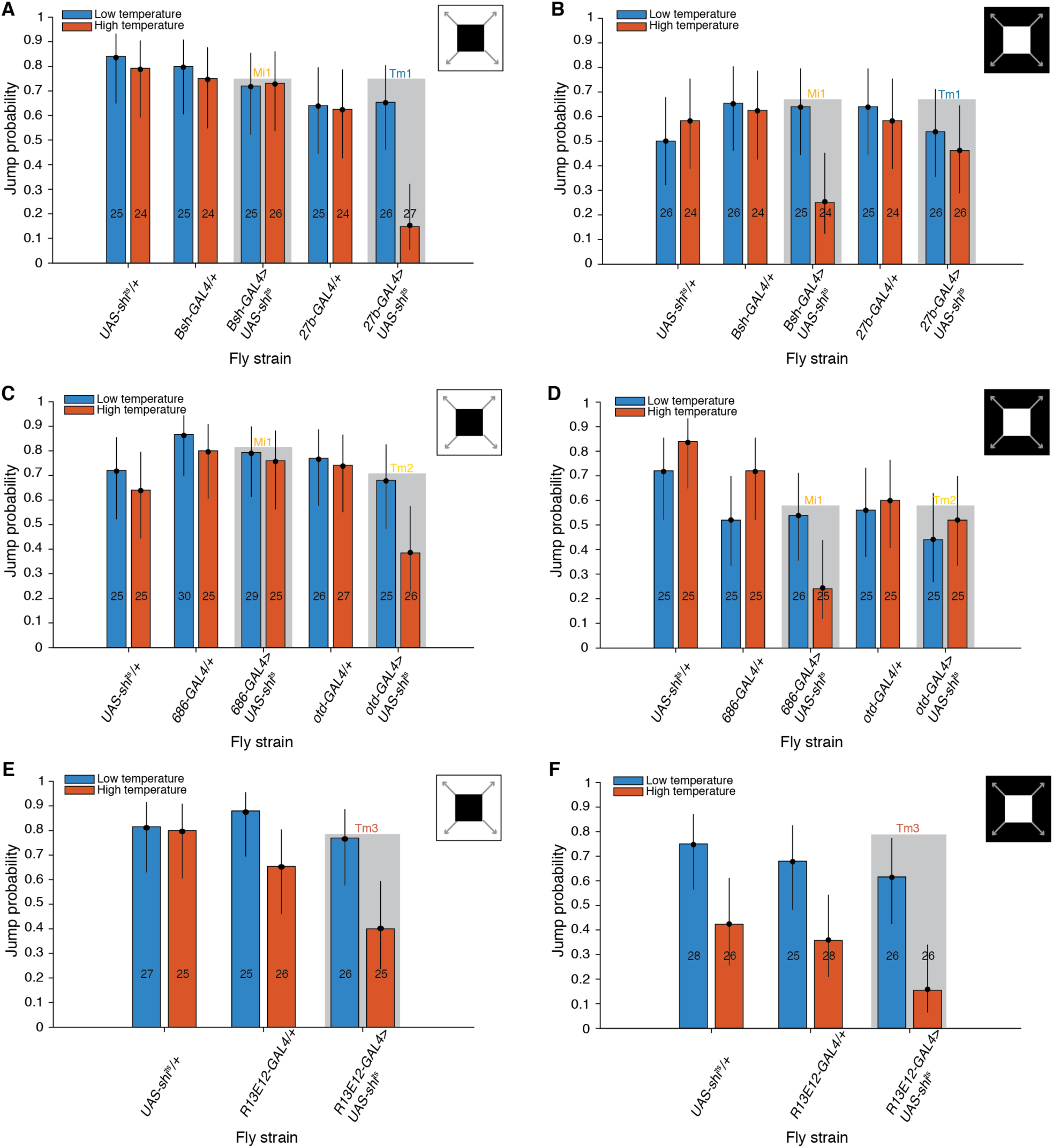
Effect of blocking medullary neurons on jump escape behavior. **A, B**: Jump probability of fly lines with either Mi1 or Tm1 neurons inactivated through *GAL4*>*UAS-shi*^*ts*^ at high temperature compared to low temperature (inside grey rectangles; ‘ON’ and ‘OFF’ looming stimuli, respectively). Control lines on *w*^*+*^ background. **C, D**: Same for lines with Mi and Tm2-blocked neurons. **E, F**: Same for lines with Tm3-blocked neurons. Plotting conventions as in Fig. 2.

**Figure 6.**
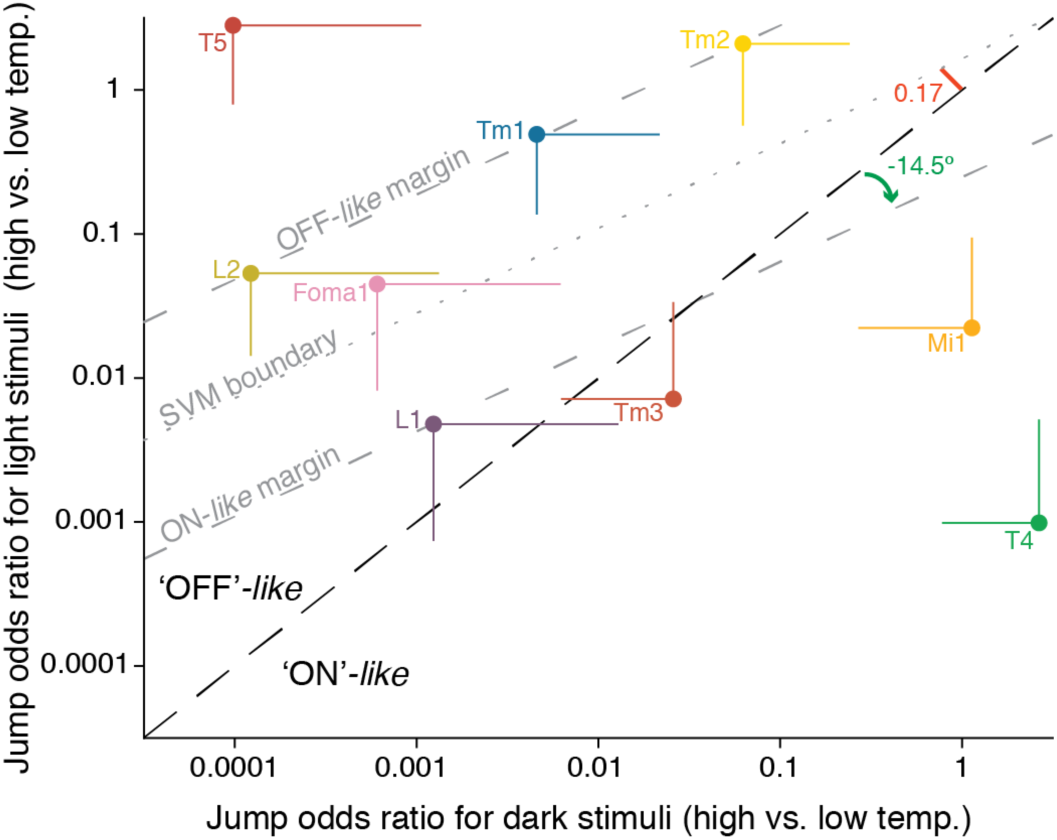
Comparison of jump odds ratios for ‘ON’ and ‘OFF’ looming stimuli across cell types. The abscissa represents the odds of jumping to a dark looming stimulus at high temperature divided by the corresponding odds at low temperature (odds ratio). The ordinate is similarly defined, but for light looming stimuli. Each point corresponds to one of the transgenic lines used in this study. Vertical and horizontal lines are standard errors. The dashed diagonal line indicates equal values for the abscissa and ordinate. The dotted grey line is the optimal boundary of the linear support vector machine classifier for ‘ON’ vs. ‘OFF’ cell types; the dashed grey lines are its margins. The angle between the boundary and the diagonal lines is indicated by a curved arrow. The distance of the boundary line to the point (1,1) on the diagonal is indicated by the thick red line. Note the double logarithmic scale.

We also applied logistic regression to compare jump probabilities across multiple fly strains at a given temperature. Note that an analysis of variance (ANOVA) cannot be applied to binary data when comparing treatment effects across multiple groups since it assumes that noise is Gaussian-distributed. We illustrate this use of logistic regression by comparing jump probabilities across multiple strains labeled 1, … *n* at the permissive temperature for *shibire*^*ts*^. One strain is selected as reference and its logit is fitted to the coefficient *η*, as above. The remaining *n* − 1 strains are assigned coefficients *α*_2_, … *α*_*n*_, as above. Strains 2, … *n* − 1 are control strains and the last strain is the one expressing *shibire*^*ts*^. We test the hypothesis that its coefficient *α*_*n*_ is equal to zero as above by using *D*_*n*−1:_ − *D*_*n*_. The value 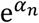 is the odds ratio of jumping for the last strain relative to that of the reference strain (log-odds ratio: *α*_*n*_).

#### Notation

In this last case, when comparing multiple strains at high temperature (ht) using logistic regression we denote by LRM_ht_[Strain 1, … Strain *n*] the logistic regression model where Strain 1 is the reference strain and Strain *n* is the test strain, for which the odds ratio is calculated relative to Strain 1 and tested for significance as described above.

#### Implementation

Logistic models were fitted by maximum likelihood using the Matlab function ‘fitglm’. The ‘Distribution’ option was set to ‘Binomial’ and the number of trials was specified with the ‘BinomialSize’ option. Instead of using *n* and *x* in the likelihood function, we used *n* and 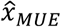, the MUE of the number successes. For each data set, 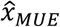 was computed by multiplying the total number of trials, *n*, and the MUE of jump probability, 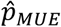, defined above. This product was rounded to the nearest tenth decimal place. In the case of a single fly strain at two temperatures (the first example above), this procedure effectively estimates the odds ratio following the method proposed by (Parzen et al. 2002). Compared to conventional logistic regression that uses directly the experimental values of *n* and *x*, this method yielded for each experiment a more conservative estimate of significance values and confidence intervals. In addition, it avoids numerical complications when the experimental jump probabilities are exactly equal to zero or one, due to the small experimental sample sizes.

#### Linear Support Vector Machine (SVM) classifier

Using the Matlab function ‘fitcsvm’ we trained a linear support vector machine to classify the jump log-odds ratios to light and dark looming stimuli for cell classes belonging to the ON and OFF pathways (Fig. 6). A linear SVM classifier is characterized by its boundary, described by the equation ***w***^*T*^***x*** + *b* = 0, where ***w*** is a two-dimensional vector perpendicular to the boundary (Cristianini and Shawe-Taylor 2000). The margin of the classifier, equal to 2/‖***w***‖, is the largest possible distance between two lines, parallel to the boundary and on either side of it, such that no member of either class lies between them.

One property of interest was the angle between the classifier boundary and the diagonal (Fig. 6, green curved arrow). A second one was the signed distance of the boundary line to the reference point (1, 1) on the diagonal, or, after taking the logarithm since we are working with log-odds ratios, the origin of the coordinate system (0, 0) (Fig. 6, red thick line). To compute the angle with respect to the diagonal, we computed the unit vector orthogonal to ***w*** which had a positive first component after rotation by −45° (thus aligning the diagonal with the horizontal axis). The second component of this rotated unit vector is equal to the sine of the angle with the diagonal, where the angle ranges from −90 to 90°. To compute the signed distance of the boundary to the point (log 1, log 1) = (0, 0), we first computed (−*b*/‖***w***‖^2^)***w***, which is the closest point on the boundary to (0, 0). The signed distance is given by the second component of this vector after it has been rotated by −45°. The sign indicates on which side of the diagonal (−*b*/‖***w***‖^2^)***w*** lies (either above and to the left, or below and to the right of it).

We computed confidence intervals for these two geometric quantities by repeating the fitting procedure 10,000 times using randomly distributed bootstrap data. For each jump odds ratio, we drew a random sample from a normal distribution with mean value equal to the experimentally determined jump log-odds ratio and variance equal to its standard error (Matlab ‘normrnd’ function). The 95% confidence intervals were obtained from the cumulative probability density function associated with the two quantities (computed with the Matlab function ‘ecdf’).

## Results

We investigated the role played by four neuron types belonging to the ON motion detection pathway of *Drosophila* in visually evoked jump escape behavior and an equal number from the OFF pathway. Both pathways are expected to provide visual input to looming-sensitive Foma1 neurons in the lobula plate of the optic lobe (Fig. 1A). Flies were placed in a funnel or holder (Fig. 1B) and presented looming stimuli simulating the approach of a dark square approaching at constant speed on a light background (Fig. 1C) or, vice-versa, a light square on a dark background. Each neuron class was inactivated by driving expression of *UAS-shibire*^*ts*^ at its permissive temperature (Methods). In *UAS-shi*^*ts*^*/+* flies, we estimated the fraction of short mode escape jumps to be 19% (n = 233 jump trials pooled between high and low temperature as their distributions were not significantly different, Kolmogorov-Smirnov test, *p* = 0.95). This value is nearly identical to that reported by von Reyn et al. (2014, their Fig. 1f).

### Similar Decrease in Looming Responses Following Inactivation of Foma1 and T4, T5

In the first experiment, we presented dark looming stimuli on a light background to six fly strains. Fig. 2A depicts their jump probability at high (permissive) and low (non-permissive) temperatures for *shibire*^*ts*^ expression, respectively. The top edge of each bar corresponds to the maximum likelihood (ML) estimate of the jump probability, i.e., the number of jumps, *x*, divided by the total number of trials, *n*. The total number of jump trials for each strain and each temperature condition is indicated either inside or above the corresponding bar. For each strain and temperature condition, a black dot indicates the median unbiased estimate (MUE) of jump probability (Methods). The MUE yielded more conservative estimates of jump probability differences than ML across strains and temperatures and was used throughout. The vertical lines above and below each MUE show 95% confidence intervals (*CI*_*95%*_) for the jump probability, computed using the Wilson score (Methods).

The largest effect of temperature on jump probability was observed for the two test fly lines, *Foma1-GAL4*>*UAS-shi*^*ts*^ and *R42F06-GAL4*>*UAS-shi*^*ts*^ (light gray shaded boxes). The first one inactivates Foma1 neurons (Fig. 2B, Supp. Fig. 1A and Supp. Movie 1) while the second one inactivates T4 and T5 neurons (Supp Fig. 1B, Supp. Movie 2). For each line, we fit logistic regression models to the high and low temperature data allowing us to quantify the strength of the temperature effect (Methods). In both cases the hypothesis that the temperature coefficient of the model, *α*_2_, was equal to zero, indicating no effect of temperature, was rejected (*p* = 2.0 10^−7^ and 8.1 10^−7^, respectively; deviance test, Methods). For each temperature, the ‘odds’ of jumping is defined as the probability of jumping, *p*, divided by the probability of not jumping, 1 − *p*. The odds ratio of jumping, computed by dividing the odds of jumping at high vs. low temperature was in both cases equal to 0.02 and deviated from the null hypothesis value of one (*CI*_*95%*_ = [0.002; 0.15] and [0.003; 0.18], respectively). An odds ratio of one indicates no change in jump odds with temperature whereas the observed odds ratio of 0.02 indicates that the odds of jumping at high temperature were fifty times less than at low temperature.

Comparing the jump probabilities at low and high temperature of the four control strains, *CantonS, UAS-shi*^*ts*^*/+, Foma1-GAL4/+* and *R42F06-GAL4/+* suggested a weak dependence on temperature (unshaded blue vs. red bars in Fig. 2A). In the last three strains, the *CI*_*95%*_ overlapped substantially at high and low temperature, showing no difference when each strain was considered in isolation. For *CantonS* flies, however, the *CI*_*95%*_ for the MUE of jump probability at high and low temperature were narrowly non-overlapping ([0.24;0.59] and [0.60;0.91], respectively). In this strain, the odds ratio of jumping at high relative to low temperature, 0.17, was 8.5 times larger than the value for the test fly lines (0.02, see above; *CI*_*95%*_ = [0.05; 0.60], *p* = 0.0037). Thus, temperature affected jumping in *CantonS* flies, but considerably less than in test flies. The weak effect of temperature on jump probability was confirmed by fitting a temperature-dependent logistic regression model to the four control strains (Methods). In this model, the temperature coefficient was different from zero (*p* = 1.6 10^−5^) and the jump odds ratio after the increase in temperature was 0.28 (*CI*_*95%*_ = [0.15; 0.51]). This decrease in the odds of jumping is again considerably smaller than what was observed in test flies.

Since temperature caused a slight decrease in jumping independent of *UAS-shi*^*ts*^ expression, we wondered if at high temperature the jump probability of *Foma1-GAL4*>*UAS-shi*^*ts*^ flies was significantly lower than that of its associated control strains: *CantonS, Foma1-GAL4/+* and *UAS-shi*^*ts*^*/+*. To address this question, we fitted a logistic regression model (LRM) to the high temperature (ht) jump probabilities of these four strains, denoted by LRM_ht_[*CantonS, Foma1-Gal4/+, UAS-shi*^*ts*^*/+, Foma1-GAL4*>*UAS-shi*^*ts*^]. We used *CantonS* as the reference strain and introduced coefficients *α*_2_, *α*_3_ and *α*_4_ for the remaining three strains (Methods). The coefficient *α*_4_ for *Foma1-GAL4*>*UAS-shi*^*ts*^ flies was different from zero (*p* = 0.0016) and the corresponding jump odds ratio of *Foma1-GAL4*>*UAS-shi*^*ts*^ to *CantonS* flies was 0.076 (*CI*_*95%*_ = [0.010; 0.559]). Hence, the odds of jumping of *Foma1-GAL4*>*UAS-shi*^*ts*^ flies were in excess of 13 times smaller than in control flies at high temperature. The same identical result held for T4T5-blocked flies, *R42F06-GAL4*>*UAS-shi*^*ts*^, when compared to their associated controls at high temperature, *CantonS, R42F06-GAL4/+* and *UAS-shi*^*ts*^*/+*.

Finally, we wondered whether differences in jump probabilities with temperature could be further resolved within the range of high or low temperatures used. For this purpose, we took advantage of the large number of trials carried out under identical conditions on *UAS-shi*^*ts*^*/+* flies depicted in Figs. 2–4. At low temperatures the data were separated in six intervals, each 0.5 °C wide. The MUE of jump probability and its 95% confidence interval were computed for each bin. Neither for dark nor for light looming stimuli was there any dependence on temperature between 19.5 and 22.5 °C (Supp. Fig. 2A, B; all *CI*_*95%*_ overlap). Similarly, at high temperature the data was split in seven intervals, each 1 °C wide, and no dependence of jump probability on temperature between 28 and 35 °C was found (Supp. Fig. 2C, D).

### Polarity Specific Decrease in Looming Responses after Inactivating T4 and T5

Next, we presented dark looming stimuli on a light background or, vice-versa, light looming stimuli on a dark background, using two strains of test flies. The first one, *R54A03-GAL4*>*UAS-shi*^*ts*^, inactivates T4 neurons (Supp. Fig. 1C, Supp. Movie 3), and the second one, *R42H07-GAL4*>*UAS-shi*^*ts*^, inactivates T5 neurons (Supp. Fig. 1D, Supp. Movie 4). Since T4 and T5 neurons are thought to belong to ON and OFF selective motion detection pathways immediately presynaptic to Foma1 neurons in the lobula (Fig. 1A), their inactivation should selectively affect responses to light and dark looming stimuli, respectively.

Inactivation of T5 neurons strongly decreased responses to dark looming stimuli (Fig. 2C, right shaded box; jump odds ratio: 0.018, *CI*_*95%*_ = [0.002; 0.138], *p* = 7.6 10^−8^), but left responses to light looming stimuli unaffected (Fig. 2D, right shaded box). Comparison of T5-blocked flies relative to control flies at high temperature produced a jump odds ratio equal to 0.015 (*CI*_*95%*_ = [0.002; 0.116], *p* = 8.8 10^−9^; LRM_ht_[*UAS-shi*^*ts*^*/+, R42H07-GAL4/+, R42H07-GAL4*>*UAS-shi*^*ts*^]). Conversely, T4 inactivation strongly decreased responses to light looming stimuli (Fig. 2D, left shaded box; jump odds ratio at high vs. low temperature: 0.049, *CI*_*95%*_ = [0.012; 0.202], *p* = 7.8 10^−7^), but left responses to dark looming stimuli unaffected (Fig. 2C, left shaded box). Further, T4-blocked flies jumped less than control flies at high temperature with a jump odds ratio equal to 0.130 (*CI*_*95%*_ = [0.042; 0.410], *p* = 2.0 10^−4^; LRM_ht_[*UAS-shi*^*ts*^*/+, R54A03-GAL4/+, R54A03-GAL4*>*UAS-shi*^*ts*^]).

In these experiments, we observed a weak difference in the odds of jumping at high vs. low temperature among control strains for dark looming stimuli (Fig 2C, those strains not enclosed in grey boxes; odds ratio: 0.46, *CI*_*95%*_ = [0.23; 0.89], *p* = 0.02), but no difference for light looming stimuli (Fig. 2D; odds ratio: 1.21, *CI*_*95%*_ = [0.63; 2.32], *p* = 0.56). Additionally, there was no significant difference in the odds of jumping for light vs. dark looming stimuli in control strains at low temperature (odds ratio: 0.54, *CI*_*95%*_ = [0.27; 1.08], *p* = 0.08).

### Lamina Monopolar Cell Inactivation Mediates Selective Decrease in Looming Responses

Next, we investigated the effect of inactivating neurons at the first stage of the ON and OFF motion detection pathways, the large monopolar cells L1 (ON) and L2 (OFF) of the lamina (Fig. 1A). We used the strain *NP6298-GAL4*>*UAS-shi*^*ts*^, which inactivates both L1 and L2 as well as *c202a-GAL4*>*UAS-shi*^*ts*^ and *21D-GAL4*>*UAS-shi*^*ts*^, which inactivate only L1 and L2, respectively (Supp. Fig. 1E-G, Supp. Movie 5-7).

We compared the simultaneous inactivation of L1 and L2 with that of Foma1. In response to dark looming stimuli, silencing both L1 and L2 decreased the jump probability (Fig. 3A, right shaded box). For these flies, the jump odds ratio at high vs. low temperature was 0.067 (*CI*_*95%*_ = [0.009; 0.494], *p* = 8.6 10^−4^). Similarly, fitting a logistic model yielded a jump odds ratio at high temperature of 0.025 for L1L2-blocked flies relative to control flies (*CI*_*95%*_ = [0.003; 0.187], *p* = 8.6 10^−7^, LRM_ht_[*UAS-shi*^*ts*^*/+, NP6298-GAL4*/+, *NP6298-GAL4*>*UAS-shi*^*ts*^]). Logistic regression revealed no dependence of the jump odds ratio on temperature among control flies (*p* = 0.20). At high temperature, the jump probability of Foma1-blocked flies also decreased, as described above, and was comparable to that of L1L2-blocked flies (Fig. 3A, left shaded box; see also Fig. 2A). The corresponding jump odds ratio was 0.040 (*CI*_*95%*_ = [0.006; 0.289], *p* = 2.1 10^−5^). At high temperature, Foma1-blocked flies jumped less than control flies with a jump odds ratio equal to 0.024 (*CI*_*95%*_ = [0.003; 0.180], *p* = 6.0 10^−7^, LRM_ht_[*UAS-shi*^*ts*^*/+, Foma1-GAL4/+, Foma1-GAL4*>*UAS-shi*^*ts*^]).

In response to light looming stimuli, the jump probability of L1L2-blocked flies was lower at high than at low temperature (Fig. 3B, right shaded box). The corresponding jump odds ratio was equal to 0.019 (*CI*_*95%*_ = [4.7 10^−4^; 0.744], *p* = 2.0 10^−4^). Similarly, at high temperature L1L2-blocked flies jumped less than control flies resulting in a jump odds ratio equal to 0.013 (*CI*_*95%*_ = [3.4 10^−4^; 0.524], *p* = 1.4 10^−5^, LRM_ht_[*UAS-shi*^*ts*^*/+, NP6298-GAL4*/+, *NP6298-GAL4*>*UAS-shi*^*ts*^]). In contrast, the jump probability of *Foma1-GAL4*>*UAS-shi*^*ts*^ flies was not different at high and low temperature (Fig. 3B, left shaded box). The jump odds ratio, 0.26, was not different from one either (*CI*_*95%*_ = [0.06; 1.10], p = 0.055). To understand this better, we computed the odds ratio of jumping to light vs. dark stimuli for *UAS-shi*^*ts*^*/+, Foma1-GAL4/+*, and *Foma1-GAL4*>*UAS-shi*^*ts*^ flies at low temperature. This revealed a significant decrease, 0.51 (*CI*_*95%*_ = [0.26; 0.97], *p* = 0.038; c.f. the three leftmost blue bars in Fig. 3A vs. those in 3B). The jump probability to light stimuli at high temperature of Foma1-blocked flies was similar to that observed earlier for dark stimuli (compare leftmost shaded red bar in Fig. 3B with those of Fig. 3A and 2A). Hence, a low baseline jump probability to light looming stimuli is likely the main culprit for the lack of significant effect at high vs. low temperature in this particular experiment.

To further investigate this issue, we also compared the jump probabilities for dark and light looming stimuli of *UAS-shi*^*ts*^*/+* control flies pooled across the experiments depicted in Figs. 2-4. At low temperature, the jump probabilities for dark looming stimuli were higher than those for light ones, *p*_*MUE*_ = 0.71 vs. 0.55 (*CI*_*95%*_ = [0.64; 0.77] and [0.47; 0.62], respectively). At high temperature, the jump probability was also slightly higher for dark vs. light looming stimuli, *p*_*MUE*_ = 0.66 vs. 0.59, but not significantly so (*CI*_*95%*_ = [0.58; 0.72] and [0.51; 0.66], respectively).

In the next experiment, we blocked L1 and L2 separately. As L1 belongs to the ON motion detection pathway, the expectation is that *c202a-GAL4*>*UAS-shi*^*ts*^ flies will jump less to light looming stimuli at the permissive temperature for *shibire*^*ts*^, whereas responses to dark looming stimuli should be unaffected. The opposite should hold for *21D-GAL4*>*UAS-shi*^*ts*^ flies since L2 belongs to the OFF pathway.

Indeed, L2-blocked flies jumped less to dark looming stimuli at high than low temperature (Fig. 3C, right shaded box; jump odds ratio: 0.020, *CI*_*95%*_ = [0.003; 0.152], *p* = 2.0 10^−7^). The jump odds ratio of L2-blocked flies relative to control flies at high temperature was 0.040 (*CI*_*95%*_ = [0.006; 0.294], *p* = 2.5 10^−5^, LRM_ht_[*UAS-shi*^*ts*^*/+, 21D-GAL4/+, 21D-GAL4*>*UAS-shi*^*ts*^]). In these experiments, we also found a significant decrease in jump probabilities of L1-blocked flies relative to their low temperature control (Fig. 3C, left shaded box; jump odds ratio: 0.055, *CI*_*95%*_ = [0.007; 0.401], *p* = 2.4 10^−4^). To further investigate the cause of the decrease, we fitted a logistic model to the low temperature (lt) data, LRM_lt_[*UAS-shi*^*ts*^*/+, c202-GAL4/+, c202-GAL4*>*UAS-shi*^*ts*^]. This revealed that *c202a-GAL4*>*UAS-shi*^*ts*^ jumped more than their controls, with a jump odds ratio of 11.2 relative to *UAS-shi*^*ts*^*/+* flies (*CI*_*95%*_ = [1.5; 83.0], *p* = 0.004). Conversely, at high temperature, the jump odds ratio of *c202a-GAL4*>*UAS-shi*^*ts*^ flies was not different from that of *UAS-shi*^*ts*^*/+* and *c202a-GAL4/+* control flies by the same logistic model analysis (*p* = 0.80). This suggests that the difference in jumping was due to a high jump probability of *c202a-GAL4*>*UAS-shi*^*ts*^ flies at low temperature rather than a selective decrease of jumping by block of L1 at high temperature.

Conversely, L1-blocked flies had a lower jump probability to light looming stimuli at high than low temperature (Fig. 3D, left shaded box; jump odds ratio: 0.098, *CI*_*95%*_ = [0.02; 0.482], *p* = 8.1 10^−4^). Similarly, in a logistic model the jump odds ratio at high temperature of the L1-blocked flies to control flies was 0.085 (*CI*_*95%*_ = [0.017; 0.423], *p* = 4.0 10^−4^, LRM_ht_[*UAS-shi*^*ts*^*/+, c202a-GAL4/+, c202a-GAL4*>*UAS-shi*^*ts*^]). In contrast, L2-blocked flies did not jump significantly less at high than low temperature (Fig. 3D, right shaded box). Yet, their jump odds ratio at high vs. low temperature, 0.28, was lower than one (*CI*_*95%*_ = [0.09; 0.86], *p* = 0.02). However, their odds ratio of jumping at high temperature relative to control flies, 0.38, was not different from one (*CI*_*95%*_ = [0.12; 1.15], *p* = 0.08, LRM_ht_[*UAS-shi*^*ts*^*/+, 21D-GAL4*>*UAS-shi*^*ts*^, *21D-GAL4*>*UAS-shi*^*ts*^]).

### Decrease in Looming Responses Following Foma1 Inactivation Confined to Central Brain

As *Foma1-GAL4/+* flies express GAL4 in the ventral nerve cord, we crossed a *tsh-GAL80* transgene (Clyne and Miesenböck 2008) into the *UAS-shi*^*ts*^ background to generate *Foma1-GAL4/tsh-GAL80; UAS-shi*^*ts*^*/+* flies that also express GAL80 in ventral nerve cord neurons. GAL80 suppresses GAL4 activity (Ma and Ptashne 1987) resulting in a substantial decrease in stained ventral nerve cord cell bodies (Supp. Fig. 3). Thus, in this line the effect of *shibire*^*ts*^ at the permissive temperature is largely restricted to central brain neurons. If Foma1 neurons in the central brain are the main drivers of jump escape behavior, we expect differences between jump probabilities at high vs. low temperature to both dark and light stimuli in double mutants. Instead, no differences would suggest a predominant role of Foma1 neurons in the ventral nerve cord.

For these experiments, we used a semi-transparent holder that allowed for better positioning of the animal in front of the visual screen and higher throughput (Fig. 1B, bottom). This led to higher jump escape probabilities in response to dark looming stimuli than with the funnel (compare Fig. 4A with Figs. 2A, 3A). Additionally, the jump probability of this line significantly decreased at high temperature (Fig. 4A, shaded box). Its jump odds ratio, 0.051, was comparable to that observed in Foma1-blocked flies (c.f. above; *CI*_*95%*_ = [0.007; 0.355], *p* = 5.6 10^−5^). The jump odds ratio of the *Foma1-GAL4/tsh-GAL80*>*UAS-shi*^*ts*^ flies was also different from that of *Foma1-GAL4/+* flies at high temperature, 0.20 (*CI*_*95%*_ = [0.057; 0.675], *p* = 5.5 10^−3^, LRM_ht_[*Foma1-GAL4/+, tsh-GAL80/+; UAS-shi*^*ts*^*/+, Foma1-GAL4/tsh-GAL80*>*UAS-shi*^*ts*^]).

In response to light looming stimuli we observed a lower jump probability than to dark looming stimuli (compare Fig. 4A and B), that was more pronounced than in earlier experiments (compare with Fig. 2A and B). The jump probability of the *Foma1-GAL4/tsh-GAL80*>*UAS-shi*^*ts*^ line decreased significantly at high temperature (Fig. 4B, shaded box), and the corresponding jump odds ratio, 0.029, was even smaller than that observed in Foma1-blocked flies (c.f. above; *CI*_*95%*_ = [0.004; 0.203], *p* = 3.4 10^−7^). The jump odds ratio of *Foma1-GAL4/tsh-GAL80* blocked flies relative to that of *Foma1-GAL4/+* flies at high temperature, 0.18, was comparable in magnitude to that for dark stimuli (0.20, see previous paragraph) but was not significantly different from one (*CI*_*95%*_ = [0.022; 1.263], *p* = 0.05, LRM_ht_[*Foma1-GAL4/+, tsh-GAL80/+; UAS-shi*^*ts*^*/+, Foma1-GAL4/tsh-GAL80*>*UAS-shi*^*ts*^]). This was presumably due to the lower baseline jump probability to light looming stimuli, which decreased our ability of distinguishing changes between the control and test strains (c.f. Discussion). Indeed, the value of the jump odds ratio at low temperature for light vs. dark looming stimuli across all strains was equal to 0.020 (*CI*_*95%*_ = [0.004; 0.090], *p* = 2.5 10^−16^).

### Moderately Selective Effect of Blocking Medullary Neurons on Looming Responses

Immediately after L1 and L2 in the lamina, the ON and OFF pathways diverge at the level of the medulla with information processed in parallel by several types of neurons converging onto T4 and T5 cells (Fig. 1A). Among those, Mi1 and Tm3 are thought to provide the most significant synaptic input to T4 (Behnia et al. 2014; Shinomiya et al. 2019; Takemura et al. 2017). Similarly, Tm1 and Tm2 are thought to be important contributors to T5 responses (Behnia et al. 2014; Shinomiya et al. 2014, 2019). We used five lines in three experiments to test the effects of inactivating these medullary neurons (Methods). For the ON pathway, we tested the lines *Bsh-GAL4*>*UAS-shi*^*ts*^ and *686-GAL4*>*UAS-shi*^*ts*^ which block Mi1 neurons (Supp. Fig. 1H, I, Supp. Movies 8, 9), as well as *R13E12-GAL4*>*UAS-shi*^*ts*^ which blocks Tm3 neurons (Supp. Fig. 1J, Supp. Movie 10). For the OFF pathway, we used the lines *27b-GAL4*>*UAS-shi*^*ts*^ which blocks Tm1 neurons and *otd-GAL4*>*UAS-shi*^*ts*^ which blocks Tm2 neurons (Supp. Fig. 1K, L, Supp. Movie 11, 12).

In the first experiment, block of Tm1 neurons resulted in a decrease in jump probability to dark looming stimuli at high vs. low temperature (Fig. 5A, right shaded box; jump odds ratio: 0.096 *CI*_*95%*_ = [0.026; 0.362], *p* = 1.4 10^−4^). At high temperature, the jump odds ratio of Tm1-blocked flies relative to control flies was 0.048 (*CI*_*95%*_ = [0.011; 0.203], *p* = 2.2 10^−6^, LRM_ht_[*UAS-shi*^*ts*^*/+, 27b-GAL4/+, 27b-GAL4*>*UAS-shi*^*ts*^]). In contrast, no change in responses to dark looming stimuli was observed when blocking Mi1 (Fig. 5A, left shaded box).

Conversely, in response to light looming stimuli the jump probability of Mi1-blocked flies was somewhat smaller at high compared to low temperature (Fig. 5B, left shaded box). However, the MUE of jump probability at high 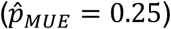 and low temperature 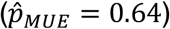 had overlapping confidence intervals (*CI*_*95%*_ = [0.12; 0.45] and [0.44; 0.80], respectively). In agreement with this weak decrease, the Mi1 jump odds ratio at high vs. low temperature was relatively large, 0.192, but different from one (*CI*_*95%*_ = [0.056; 0.655], *p* = 0.006). A similar result held for the jump odds ratio at high temperature of Mi1-blocked flies relative to control flies, which was equal to 0.24 (*CI*_*95%*_ = [0.071; 0.829], *p* = 0.02, LRM_ht_[*UAS-shi*^*ts*^*/+, Bsh-GAL4/+, Bsh-GAL4*>*UAS-shi*^*ts*^]). No change in jump probability in response to light looming stimuli was detected for Tm1-blocked flies (Fig. 5B, right shaded box).

In the second experiment, block of Tm2 neurons led to a slight decrease in jump probability to dark looming stimuli at high temperature that was not significant (Fig. 5C, right shaded box). Correspondingly, the jump odds ratio at high relative to low temperature of Tm2-blocked flies was relatively high, 0.30 (*CI*_*95%*_ = [0.095; 0.947], *p* = 0.036). At high temperatures, the jump odds ratio of Tm2-blocked flies relative to control flies was not different from one (0.352, *CI*_*95%*_ = [0.113; 1.095], *p* = 0.066, LRM_ht_[*UAS-shi*^*ts*^*/+, otd-GAL4/+, otd-GAL4*>*UAS-shi*^*ts*^]). As in the previous experiment, blocking Mi1 neurons did not affect jumping responses to dark stimuli (Fig. 5C, left shaded box).

In response to light looming stimuli, no change in jump probability with temperature was observed for Tm2-blocked flies (Fig. 5D, right shaded box). Block of Mi1 neurons resulted in a decrease in jump probability at high temperature that was not significant (Fig. 5D, left shaded box). The odds ratio of jumping at high vs. low temperature was fairly large, 0.28 (*CI*_*95%*_ = [0.084; 0.912], *p* = 0.030). The odds ratio of jumping at high temperature of Mi1-blocked flies relative to control flies was however significantly lower, 0.063 (*CI*_*95%*_ = [0.016; 0.256], *p* = 1.4 10^−5^, LRM[*UAS-shi*^*ts*^*/+, 686-GAL4/+, 686-GAL4*>*UAS-shi*^*ts*^]).

In the third experiment, Tm3-blocked flies had a decreased probability of jumping in response to dark looming stimuli at high vs. low temperature, 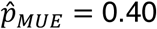 vs. 0.77 (Fig. 5E, shaded box). This decrease was, however, not significant as expected for a neuron belonging to the ON pathway (*CI*_*95%*_ = [0.24; 0.59] vs. [0.58; 0.89]). Accordingly, the jump odds ratio after the temperature increase was relatively high, 0.204 (*CI*_*95%*_ = [0.061; 0.685], *p* = 0.007). Similarly, the jump odds ratio at high temperature of Tm3-blocked flies relative to control flies was equal to 0.171 (*CI*_*95%*_ = [0.048; 0.602], *p* = 0.004, LRM_ht_[*UAS-shi*^*ts*^*/+, R13E12-GAL4/+, R13E12-GAL4*>*UAS-shi*^*ts*^]).

In contrast, in response to light looming stimuli the jump probability of Tm3-blocked flies was lower at high vs. low temperature (Fig. 5F, shaded box). The jump odds ratio was moderately decreased, 0.117 (*CI*_*95%*_ = [0.031; 0.437], *p* = 5.1 10^−4^) but the jump odds ratio of Tm3-blocked flies at high temperature relative to control flies was fairly high, 0.255 (*CI*_*95%*_ = [0.069; 0.947], *p* = 0.033, LRM_ht_[*UAS-shi*^*ts*^*/+, 13E12-GAL4/+, 13E12-GAL4*>*UAS-shi*^*ts*^]). This appeared due to a weak decrease in jump probability at high temperature. Indeed, the jump odds ratio at high vs. low temperature of control flies, *UAS-shi*^*ts*^*/+* and *13E12-GAL4/+*, showed a moderate decrease of 0.256 (*CI*_*95%*_ = [0.114; 0.574], *p* = 6.6 10^−4^).

### Jump Odds Ratios Recapitulate the Dichotomy Between ON and OFF Pathways

To jointly characterize the effect of inactivating different cell types on jump escape behavior, we plotted the log-odds ratios of jumping in response to light looming stimuli at high vs. low temperature against the log-odds ratios for dark looming stimuli (Fig. 6). Inactivating cell types that lie below the diagonal more strongly affects the odds of jumping to light rather than dark stimuli. Vice-versa, inactivating those that lie above the diagonal more strongly affects the odds of jumping to dark rather than light looming stimuli (‘ON-like’ and ‘OFF-like’, respectively). Notably, all neurons belonging to the OFF pathway clustered above the diagonal (L2, Tm1, Tm2, T5; Fig. 1A). Conversely, the cell types Tm3, Mi1 and T4 that belong to the ON pathway lied below the diagonal. L1 was the only ON pathway neuron found above the diagonal, but less than one standard error away from it along both axes (Fig. 6, horizontal and vertical lines stemming from each data point). This could be traced back to the high jump probability of L1 control flies at low temperature already discussed in the context of Fig. 3C. To discriminate between ON and OFF cells, we used a linear support vector machine (SVM) classifier. The linear SVM classifier maximizes the distance, called the ‘margin’, of two parallel lines between which no member of either class lies (Fig. 6, dashed grey margin lines; Methods). The corresponding classifier boundary line, located half-way from either margin line, perfectly separated the two cell classes (Fig. 6, dotted grey line). The Foma1 data point lied within the margins, but 42% closer to the ‘OFF-like’ than the ‘ON-like’ margin. The angle of the classifier boundary line relative to the diagonal line was −14.5° (Fig. 6, curved arrow; Methods). After accounting for the uncertainty in jump odds ratios (Fig. 6, illustrated by standard errors), we estimated the angle mean value to be −20.3° and its 95% confidence interval [-46.0°; 9.1°] (see Methods), which was thus not different from 0°. We also computed the signed distance of the boundary to the diagonal at the reference point (1,1). It amounted to 0.17 log units (Fig. 6, thick red line; Methods). Accounting for the uncertainty in jump odds ratios, yielded a mean distance of −0.22 log units. As the 95% confidence interval was [–1.07; 0.91] log units, the distance of the boundary from the diagonal was not significantly different from zero either.

## Discussion

We investigated the role of nine cell classes in the generation of *Drosophila* jump escape behaviors evoked by dark and light looming stimuli in acute silencing experiments. Four of these, L1, Mi1, Tm3 and T4 belong to the ON motion detection pathway, and another four belong to the OFF pathway, L2, Tm1, Tm2 and T5 (Fig. 1A). The last one, Foma1, comprises lobula plate output neurons sensitive to both light and dark looming stimuli. Under identical behavioral conditions, the silencing of Foma1 closely resembled that of T4 neurons in response to light stimuli and of T5 neurons to dark stimuli. Similarly, experiments silencing Foma1 or L1 and L2, as well as experiments silencing either L1 or L2 separately, showed a tight correspondence between the behavioral effects of silencing L1 and Foma1 for light looming stimuli and, correspondingly, between silencing L2 and Foma1 for dark looming stimuli. Taken together, these results suggest that the ON pathway, initiating with L1 in the lamina and terminating with T4 in the lobula plate, provides the main synaptic input to Foma1 lobula plate neurons for light looming stimuli. Conversely, the OFF pathway starting with L2 and ending with T5 likely fulfills the same role for dark looming stimuli. These conclusions are in agreement with behavioral observations of visually guided collision avoidance in flight (Schilling and Borst 2015).

In flies with GAL80 suppressed activation of GAL4 in the ventral nerve cord, we observed a larger difference between the jump probabilities to dark and light stimuli compared with flies where Foma1 neurons were inactivated across the entire nervous system (compare leftmost shaded red bars in Figs. 4A, B with those in Fig. 3A, B). One key difference between these experiments was the use of a holder (in Fig. 4) aligning the right eye of the flies with the screen rather than a funnel (in Fig. 3) allowing flies to emerge at different orientations relative to the screen (see also Fig. 1B). Indeed, a close inspection of the filmed behavior revealed that in the GAL80 suppression experiments of Fig. 4 flies were consistently less eager to step on the holder’s platform when their right eye directly faced a completely dark screen as opposed to one with a light background. This behavior was observed irrespective of the fly strain or experimental temperature. Consequently, the use of the holder and its positioning relative to the screen likely accentuated the difference in baseline jump probabilities to dark and light looming stimuli. Yet, the jump probabilities to both dark and light stimuli were significantly decreased at the permissive temperature in these *Foma1-GAL4/tsh-GAL80*>*UAS-shi*^*ts*^ mutants, with jump odds ratios comparable or even smaller than those of Foma1 flies for which *shibire*^*ts*^ inactivation is not suppressed in the ventral nerve cord. Thus, we conclude that Foma1 neurons in the central nervous system are sufficient to explain our results on jump escape behavior to light and dark looming stimuli. However, given the difference in baseline jump probability observed in the ventral nerve cord suppressed and unsuppressed experiments of Fig. 3 and 4, we cannot rule out that Foma1 neurons in the ventral nerve cord modulate overall jump probability to light and dark looming stimuli.

Just as the Foma1 line drives GAL4 expression in neurons outside the optic lobe, the other lines used in this study also induce GAL4 expression in varied patterns outside the optic lobes. Yet, the segregation of neurons into ON and OFF pathways has been observed up to now only at early stages of visual processing in the optic lobes (Fig. 1A). It thus appears unlikely that neurons outside the optic lobe could be the main determinants of our behavioral results because of the tight correlation observed in each experiment between jump behavior after *shibire*^*ts*^ inactivation and looming stimulus polarity. Hence, either ON/OFF segregation persists beyond the well documented circuits in the optic lobes for which there is presently no evidence, or more plausibly, the specific optic lobe neurons belonging to the ON/OFF pathways targeted by our driver lines are responsible for the behavioral dichotomy based on stimulus polarity documented here.

At the level of the medulla, the effect of inactivating neurons belonging to either pathway resulted in stimulus polarity-specific changes in jump escape behavior that were less clear-cut than those observed when silencing the initial or final elements of each pathway. This likely resulted from silencing only a subset of the neurons that receive input from L1 and L2 in the medulla and provide input to T4 and T5 (Shinomiya et al. 2019). Additionally, the ON and OFF motion pathways do not appear to be completely segregated at the level of the medulla (Fisher et al. 2015; Shinomiya et al. 2019).

Using logistic regression and a SVM classifier, we quantified the joint effect of silencing each neuronal class on jump escape behavior to light and dark looming stimuli. This revealed that we could perfectly segregate neurons in ‘ON-like’ and ‘OFF-like’ categories based solely on behavioral jump odds ratios (Fig. 6). Additionally, the boundary between the two categories was close and not significantly different from the diagonal. This outcome is exactly what is expected if neurons belonging to the anatomical ON pathway affect more responses to light looming stimuli and vice-versa for OFF neurons. Thus, these findings further suggest a tight link between ON and OFF motion sensitive neuronal pathways and the generation of escape behaviors to light and dark looming stimuli. We interpret the location of Foma1 neurons closer to ‘OFF-like’ cells in the log-odds ratios space of Fig. 6 as an indication of the general bias of the visual system towards generating escape behaviors to dark rather than light stimuli as reported above in several individual experiments. Biases for dark vs. light contrasts have been documented both in vertebrate and invertebrate visual systems (e.g., Chen et al. 2019; Leonhardt et al. 2016; Mazade et al. 2019).

Escape behaviors to looming stimuli have been particularly well investigated in two other family of arthropods: grasshoppers and crabs. In contrast to *Drosophila*, the visual inputs to looming detection neurons in these animals are not thought to be directionally selective (Jones and Gabbiani 2010, 2012; Medan et al. 2007; Oliva and Tomsic 2016). Yet, the computations carried out in *Drosophila*, grasshoppers, and crabs appear very similar at the behavioral level (Fotowat et al. 2011; Oliva and Tomsic 2012; von Reyn et al. 2014). The biophysics of looming sensitive neurons has long been investigated in grasshoppers and is increasingly well understood in *Drosophila* (Ache et al. 2018; von Reyn et al. 2017). Thus, grasshoppers and *Drosophila* present attractive models to compare and elucidate the relative advantages of harnessing direction-selective and non-direction selective neural circuits for jump escape and how their differences may impact behavior.

## Acknowledgements

Stocks obtained from the Bloomington Drosophila Stock Center (NIH P40OD018537) were used in this study. The anti-Dlg monoclonal antibody, developed by Dr. Goodman, was obtained from the Developmental Studies Hybridoma Bank, created by the NICHD of the NIH and maintained at The University of Iowa, Department of Biology, Iowa City, IA 52242. We thank Drs. Clandinin, Venken, and Miesenböck for generously sharing stocks.

## Data availability

Data and Matlab code reproducing the main figures will be made available on Mendeley.

## Supplementary Material

**Supplementary Figure 1.**
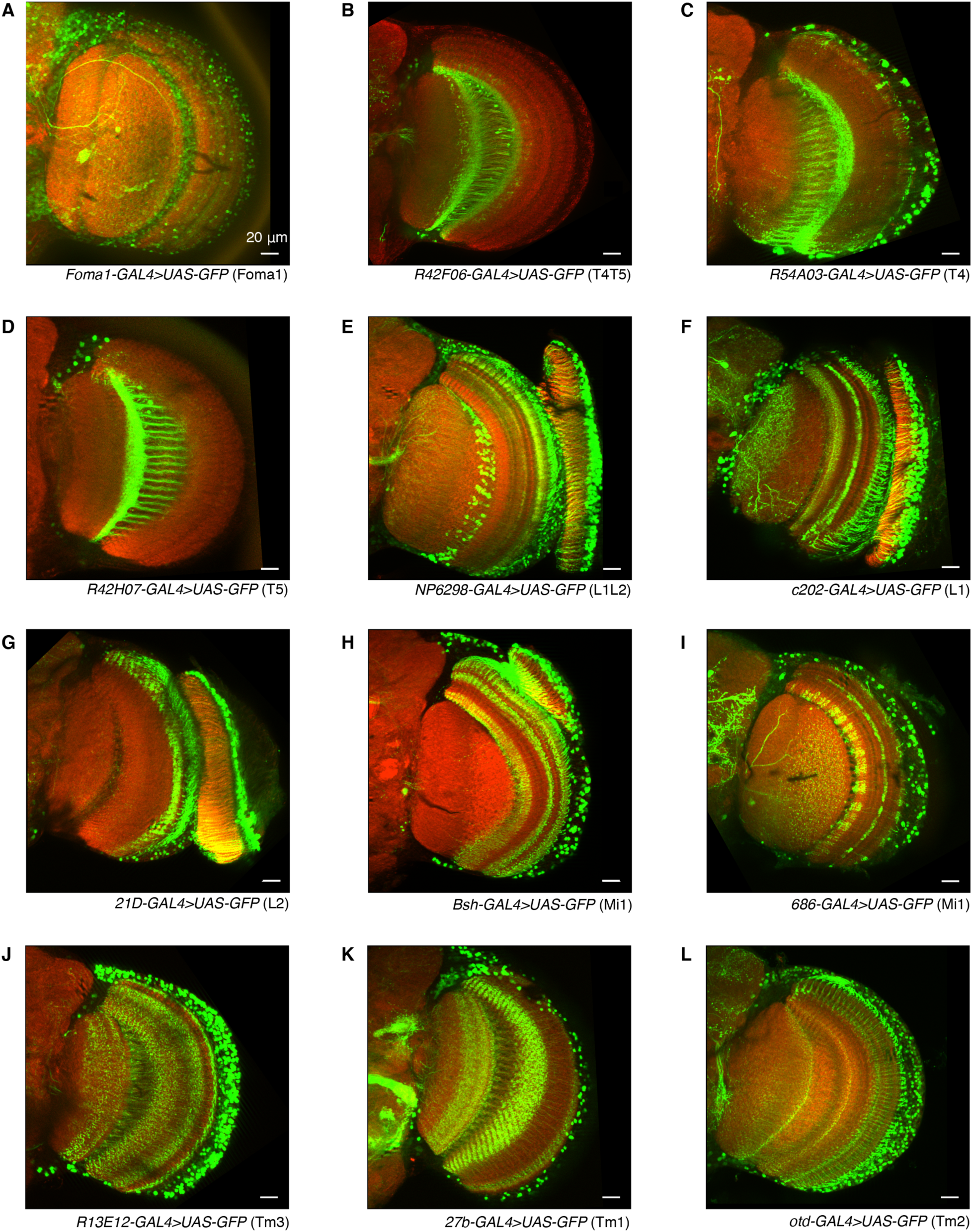
Staining patterns in the optic lobes of the fly transgenic driver lines used in this study. These are *Foma1-GAL4*>*UAS-GFP* (Foma1, **A**), *R42F06-GAL4*>*UAS-GFP* (T4 and T5, **B**), *R54A03-GAL4*>*UAS-GFP* (T4, **C**), *R42H07-GAL4*>*UAS-GFP* (T5, **D**), *NP6298-GAL4*>*UAS-GFP* (L1 and L2, **E**), *c202-GAL4*>*UAS-GFP* (L1, **F**), *21D-GAL4*>*UAS-GFP* (L2, **G**), *Bsh-GAL4*>*UAS-GFP* (Mi1, **H**), *686-GAL4*>*UAS-GFP* (Mi1, **I**), *R13E12-GAL4*>*UAS-GFP* (Tm3, **J**), *27b-GAL4*>*UAS-GFP* (Tm1, **K**), and *otd-GAL4*>*UAS-GFP* (Tm2, **L**). Scale bars: 20 *µ*m.

**Supplementary Figure 2.**
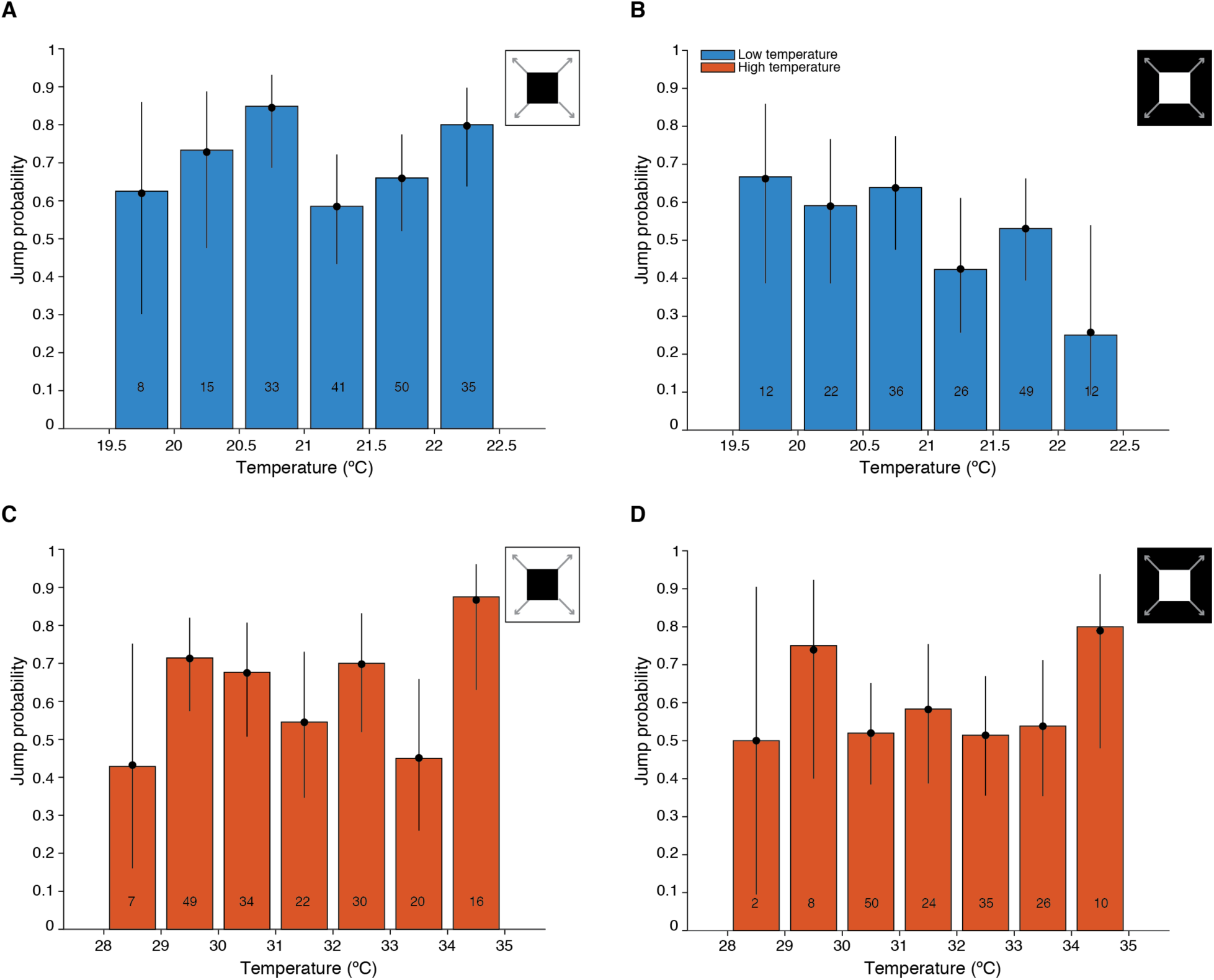
Independence of jump escape behavior on fine scale temperature changes. **A, B:** Jump probability of *UAS-shi*^*ts*^*/+* flies at low temperature, broken down in 0.5 °C bins (‘ON’ and ‘OFF’ looming stimuli, respectively). **C, D:** Jump probability of *UAS-shi*^*ts*^*/+* flies at high temperature, broken down in 1 °C bins (‘ON’ and ‘OFF’ looming stimuli, respectively.

**Supplementary Figure 3.**
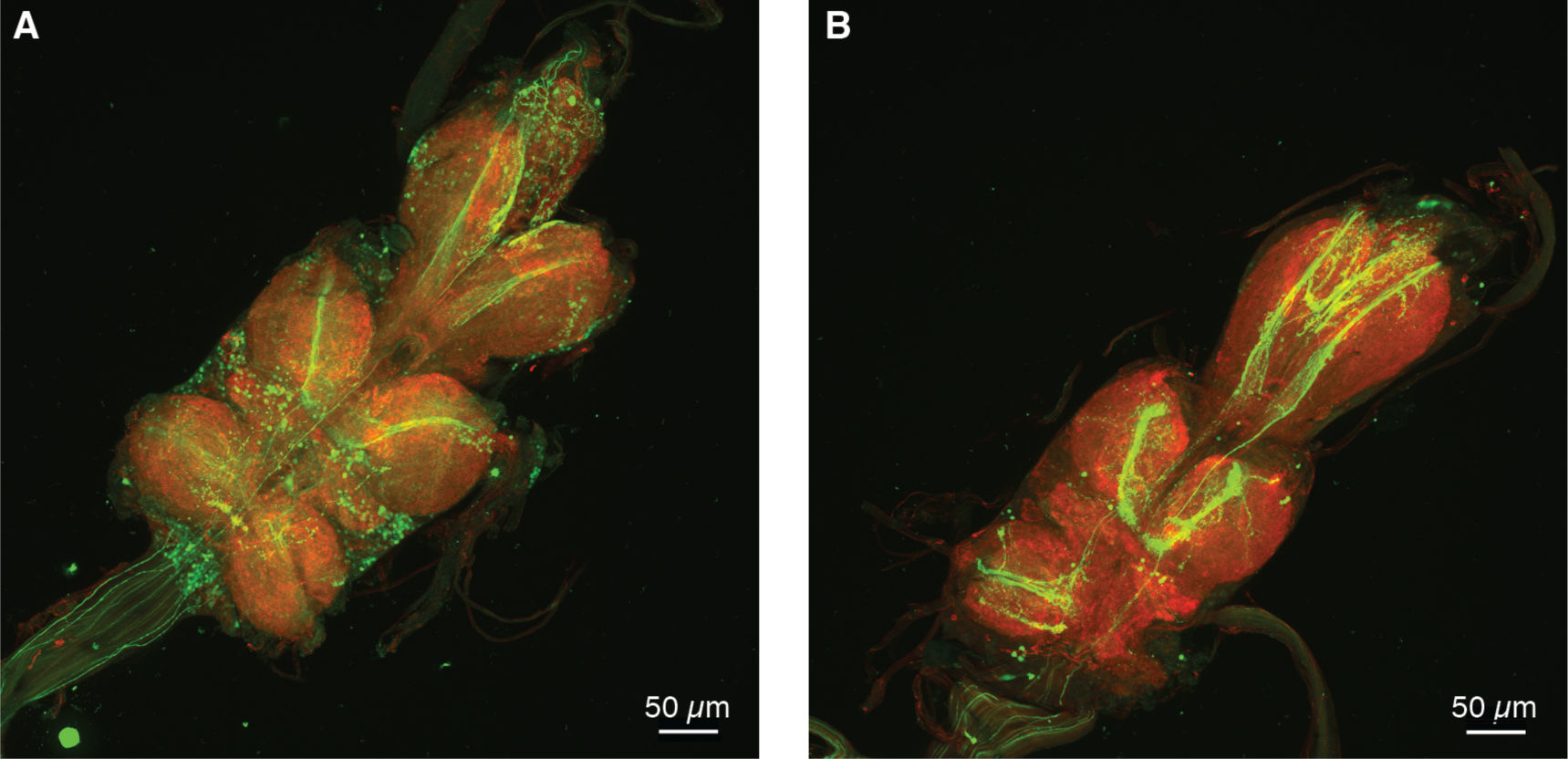
Staining patterns of *Foma1-GAL4*>*UAS-GFP*. (**A**) and *Foma1-GAL4/tsh-GAL80*>*UAS-GFP* (**B**) in the ventral nerve cord. Note the strong decrease in neuronal cell body stained in the *tsh-GAL80*-expressing line. Scale bars: 50 *µ*m.

**Supplementary Figure 4.**
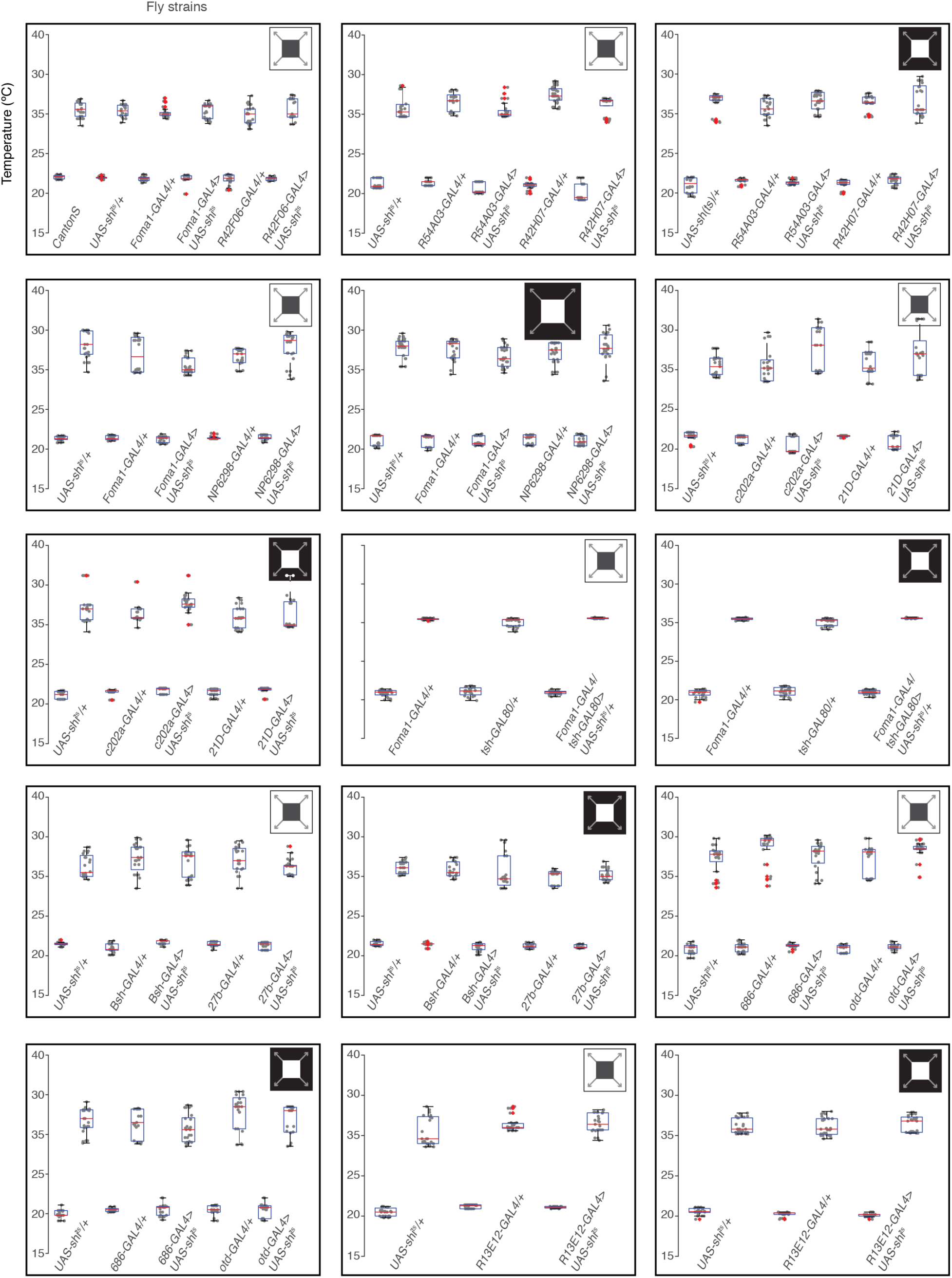
Temperature data for each experiment. Each plot reports the temperature recorded for each trial at high and low temperature. The experiments are arranged from left to right and top to bottom in the same order of presentation as in the main text. For all strains (indicated below) and temperature conditions (low followed by high), each grey dot represents one jump trial (the position of grey dots along the x-axis was randomized for increased visibility). The red central line represents the median of the temperature distribution and the blue bottom and top edges of each box the 25^th^ and 75^th^ percentiles. The whiskers extend to the most extreme points not considered outliers and the red crosses represent outliers.

**Supplementary Movies 1-12.** Animation of slices through the optic lobes of the same *Drosophila* lines illustrated in Supp. Fig. 1A-L (Supp. Movies 1-12, respectively). Scale bars: 20 *µ*m.

